# Causal dependencies between frontal and temporal lobe regions underlying word search and retrieval

**DOI:** 10.64898/2026.05.20.726706

**Authors:** Benedikt Winzer, William Burns, Rishika Chikoti, Emma Strawderman, Steven P. Meyers, Kevin A. Walter, Webster H. Pilcher, Madalina E. Tivarus, Bradford Z. Mahon, Frank E. Garcea

## Abstract

Verbal fluency is a behavioral task that requires the generation of words from a semantic category (category fluency) or words beginning with a specific letter (letter fluency). Although word production engages a frontal-temporal-parietal network, no studies have tested how lesions to temporal and parietal lobe areas that represent semantic and phonological knowledge dampen neural responses in the left pars triangularis and the left pars opercularis, two adjacent regions in the left inferior frontal gyrus implicated in word search and retrieval. Here, 52 patients with temporal lobe lesions underwent clinical functional MRI while performing the category and letter fluency tasks. We investigated where lesion presence was inversely related to the magnitude of task-specific neural responses in pars triangularis and pars opercularis using a technique referred to as voxel-based lesion activity mapping (VLAM). We found that lesions to the left anterior superior temporal gyrus, left temporal pole, left hippocampus, left insula, and underlying inferior fronto-occipital fasciculus were associated with reduced neural responses in the left pars triangularis during the category fluency task. Lesion damage to the right hippocampus was associated with reduced neural responses in the left pars opercularis during category fluency. By contrast, lesions to the left posterior superior temporal gyrus, left supramarginal gyrus, left parietal operculum, and the inferior fronto-occipital fasciculus and left arcuate fasciculus were associated with reduced neural responses in the left pars triangularis and the left pars opercularis during the letter fluency task. These results suggest that anatomically dissociable brain networks interact with the left inferior frontal gyrus when different search strategies constrain the retrieval of word representations.

## Introduction

Fluent word production depends upon coordinated retrieval processes across separable stages, including semantics, lexical, phonological, and phono-articulatory levels of representation. Verbal fluency is a paradigm in which participants are given 1-2 minutes to produce words from a semantic (e.g., animals, fruits, clothing; hereafter, category fluency) or phonological category (e.g., words beginning with A, F, or S; hereafter, letter fluency). Producing words under these constraints engages semantic memory, retrieval of lexical and phonological word knowledge, working memory, self-monitoring, and set switching to generate words while avoiding repetitions (Martin et al., 1994; Moscovitch, 1995; Rende et al., 2002; Rosen & Engle, 1997; Troyer et al., 1997). Voxel-based Lesion-Symptom Mapping (VLSM) studies (Baldo et al., 2006; Biesbroek et al., 2016; Li et al., 2017; Stuss & Levine, 2002; Thye et al., 2021) and functional Magnetic Resonance Imaging (fMRI) investigations (Birn et al., 2010; Katzev et al., 2013; Noppeney et al., 2004; Perani et al., 2003) converge in demonstrating that word production during verbal fluency recruits a distributed network that includes the inferior frontal gyrus, the anterior temporal lobe, and temporo-parietal regions. Here, we combine VLSM with fMRI to investigate functional interactions between the left inferior frontal gyrus and left temporo-parietal areas involved in word search and retrieval as patients with lesions take part in the category and letter fluency tasks.

Word production in the verbal fluency task is executively demanding, as it requires endogenous retrieval of semantically or phonologically related words. fMRI studies have shown there are increased neural responses in the left inferior frontal gyrus when task demands are more difficult (Fedorenko et al., 2012) or when they require controlled retrieval of semantic representations (Badre & Wagner, 2007; Jackson, 2021; Ralph et al., 2017; Thompson-Schill et al., 2005). Similar neural effects have been observed in the left inferior frontal gyrus when participants engage in verbal fluency tasks (Birn et al., 2010; Hirshorn & Thompson-Schill, 2006; Katzev et al., 2013), consistent with the idea that verbal fluency, as a task, is dependent upon executive control systems when retrieving words during language production.

Neuropsychological studies provide causal evidence demonstrating that partially distinct temporal lobe regions underlie the retrieval of words representations at different stages of processing (for discussion, see Dell et al., 2013; Hickok & Poeppel, 2007; Piai & Eikelboom, 2023). For example, lesions to anterior portions of the left superior and middle temporal gyri are associated with semantic errors in picture naming (e.g., saying ‘cow’ for ‘horse’; Baldo et al., 2013; Cloutman et al., 2009; Mirman, Zhang, et al., 2015; Schwartz et al., 2009), whereas lesions to the left supramarginal gyrus are associated with phonological errors in picture naming (e.g., saying ‘horsh’ for ‘horse’; Schwartz et al., 2012), deficits in verbal working memory (Buchsbaum et al., 2011), and difficulty with speech repetition (Fridriksson et al., 2010).

Although VLSM studies have identified common lesion sites associated with performance in the category and letter fluency tasks (Biesbroek et al., 2021; Chouiter et al., 2016; Thye et al., 2021), distinct areas have also been linked to performance across the tasks. For instance, lesions to the left anterior superior and middle temporal gyri (Baldo et al., 2006; Zigiotto et al., 2022), left hippocampus (Biesbroek et al., 2016), and underlying left inferior fronto-occipital fasciculus (IFOF; Almairac et al., 2015) have been associated with fewer words produced in the category fluency task but not the letter fluency task. By contrast, lesions to the left posterior inferior frontal gyrus and left precentral gyrus (Baldo et al., 2006; Schmidt et al., 2019), the left supramarginal gyrus (Chouiter et al., 2016), and the left arcuate fasciculus (Blecher et al., 2019) have been associated with fewer words produced in the letter fluency task but not the category fluency task. These findings suggest that the two tasks rely on dissociable neural pathways that support different retrieval strategies (Katzev et al., 2013; Schmidt et al., 2017). In the category fluency task, activation spreads among semantically related concepts cued by the instruction, requiring participants to switch among semantic subcategories (Gruenewald & Lockhead, 1980; Troyer et al., 1997). In the letter fluency task, words may be retrieved via phonologically-constrained search strategies, such as syllabification of the initial letters (Mummery et al., 1996), though semantic memory may provide a scaffold to structure the retrieval of phonologically related concepts (e.g., see Schwartz et al., 2003).

Our proposal is that activation in the left inferior frontal gyrus represents a summation of endogenous control processes that guide the retrieval of conceptual representations for lexical selection. The retrieval process depends, by hypothesis, on dynamic interactions with task-relevant representations that are the target of the search. During category fluency, when searching occurs among semantically related concepts, we hypothesize that functional interactions between the left inferior frontal gyrus and anterior portions of the left middle and superior temporal gyri refine the retrieval of semantically related concepts for further processing by the lexical system. During letter fluency, we hypothesize that functional interactions among the left inferior gyrus, the left posterior superior temporal gyrus, and the left supramarginal gyrus refine the search of phonologically related concepts for processing by the lexical system. This neuro-cognitive framework makes the prediction that: 1) Participants with lesions to anterior portions of the left middle and superior temporal gyri will show reduced neural responses in the left inferior frontal gyrus during category fluency, reflecting disrupted coordination between the regions mediating retrieval of semantically related concepts; and 2) Participants with lesions to the left posterior superior temporal gyrus and supramarginal gyrus will show reduced neural responses in the left inferior frontal gyrus, reflecting disrupted coordination between the regions mediating retrieval of phonologically related concepts.

To test these predictions, we analyzed category and letter fluency fMRI data collected in 52 individuals with acquired brain lesions concentrated in the left and right temporal lobe, and the white matter undercutting the left inferior parietal lobule. Lesions were normalized to a common stereotactic space, allowing us to test if distinct temporal and parietal lesion sites disrupt neural responses in the left inferior frontal gyrus in a task-specific manner. The analysis of neural responses was restricted to the left pars triangularis and the left pars opercularis given that both regions have been identified in fMRI studies using the verbal fluency task (e.g., see Katzev et al., 2013). We report findings separately using each region-of-interest (ROI) for hypothesis testing. In each analysis, neural responses in from the second verbal fluency task are entered as a covariate (e.g., to control for letter fluency when inspecting the lesion sites associated with reduced neural responses in the category fluency task, and vice versa).

Deterministic fiber tracking in neurotypical participants then tested whether distinct white matter fibers mediate structural connectivity between the left inferior frontal gyrus region and the lesion sites identified in the category and letter fluency analyses. Building from our initial set of predictions, and using the lesion sites identified in the analyses described above as ROIs, we hypothesized that 3) Anterior portions of the left middle and superior temporal gyrus are structurally connected to the left inferior frontal gyrus via the left IFOF and/or uncinate fasciculus; and 4) The left superior temporal gyrus and left supramarginal gyrus are structurally connected to the left inferior frontal gyrus via the left IFOF and/or arcuate fasciculus. If these hypotheses are supported, it suggests that dissociable networks underlie the retrieval of semantically and phonologically related concepts to constrain the selection of lexical representations for word production.

## Methods

### Participants

Over a six-year period, 206 individuals underwent clinical fMRI as part of their pre-operative neurosurgical clinical care at the University of Rochester Medical Center. MRI data were exported from the medical imaging archive system and identifying information was removed from the DICOM headers. Data analysis was restricted to participants who met the following inclusion criteria: (1) Were between the ages of 18 and 80 at the time of the scan, (2) Had no prior history of brain surgery, and (3) Had an identifiable lesion on the T1 image. Fifty-two of the 206 participants who met these inclusion criteria completed the category and letter fluency tasks. Of those participants, 37 had left hemisphere lesions and 15 had right hemisphere lesions (20 females; mean age, 42.6 y; SD 15.5 y). All analyses were carried out in this group (see Figure 1 for a lesion overlap map and Supplemental Table 1 for demographic variables). This retrospective research study was approved by the University of Rochester Research Subjects Review Board.

**Figure 1.**
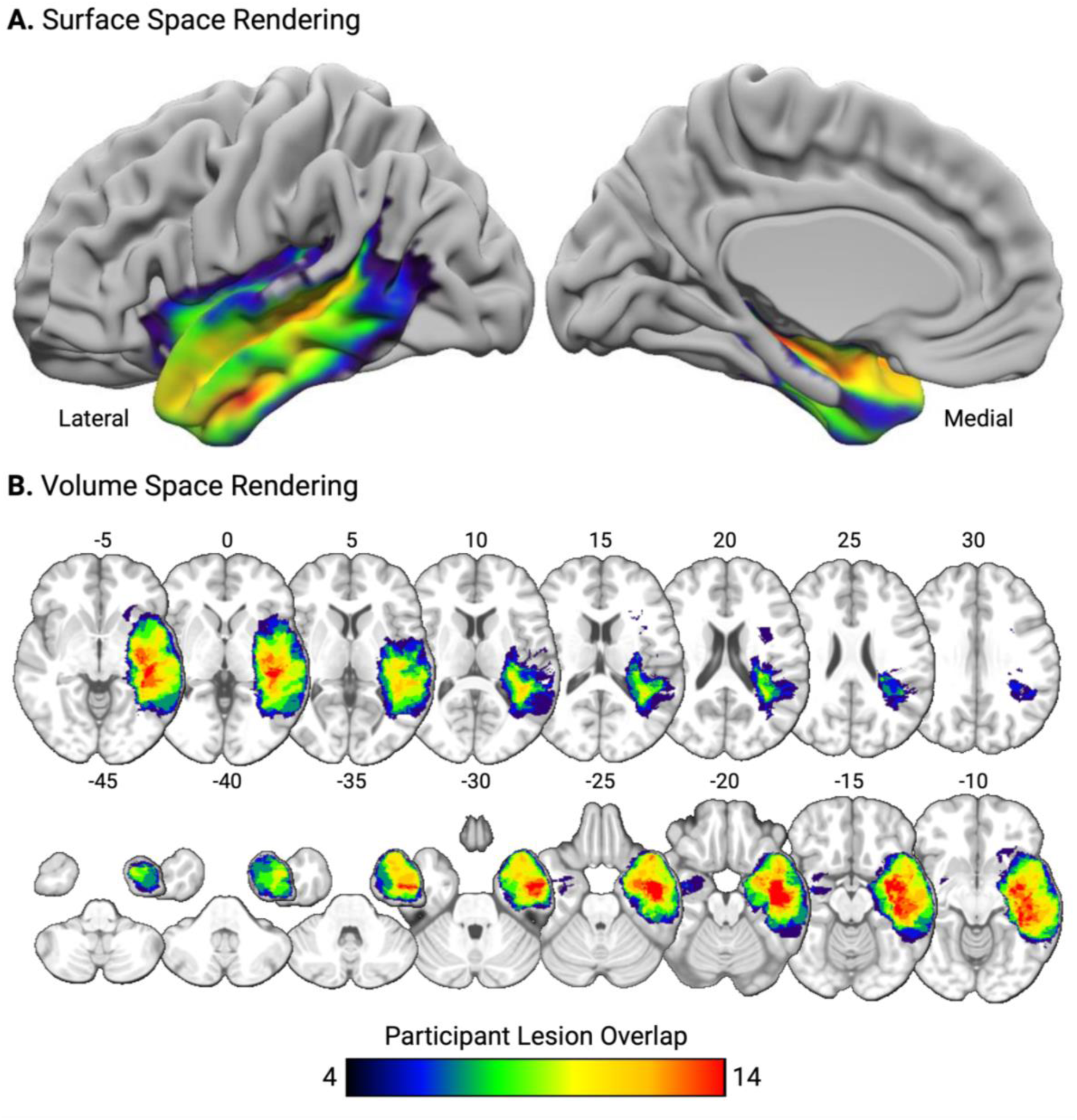
Voxelwise lesion overlap map among the 52 participants projected (A) on a surface representation of the ICBM152 brain and (B) in volume space in radiological convention. MNI Z coordinates are listed above each axial slice.

Fifty-four neurotypical adults participated in an fMRI and diffusion MRI protocol as part of a separate project investigating object representations in the ventral visual pathway (29 females; mean age, 22.27 y; SD 6.03 y; see Chen et al., 2016; Erdogan et al., 2016; Garcea et al., 2016; Kristensen et al., 2016). Scanning took place at the Center for Advanced Brain Imaging and Neurophysiology at the University of Rochester. Neurotypical participants had no history of psychiatric illness or neurologic injury and were right-hand dominant. Participants had normal or corrected-to normal vision, took part in the study in exchange for payment, and gave written informed consent in accordance with the University of Rochester Research Subjects Review Board. For the purpose of this project, we restricted our analysis to the diffusion MRI data.

### Experimental Procedure

Each participant with a brain lesion underwent a high resolution T1 anatomical scan, followed by the letter fluency task and the category fluency task. Sensory and motor mapping tasks were administered after these fluency tasks but are not analyzed herein, as they are not germane to the focus of this investigation. Participants were familiarized with the stimuli and practiced each task prior to the scan to ensure they could sustain attention and follow task instruction. The category fluency task cued the subvocal production of words from different semantic categories. Each 30-second fluency block began with a visual stimulus cueing participants to silently produce as many words as possible belonging to a specified semantic category. The first semantic category was animals, followed by foods, and furniture. The letter fluency task cued the subvocal production of words starting with different letters. Each 30-second fluency block began with a visual stimulus cueing participants to silently produce as many words as possible starting with a specified letter. The first letter category was A, followed by F, and S. Across both tasks, the category or letter cue was presented for the full duration of the fluency block, followed by 30-second fixation blocks. Each run consisted of 4 fixation blocks interspersed with 3 fluency blocks.

One run of each task was collected, with the caveat that an additional run was performed if the participant had difficulty performing the task (e.g., if they did not attend to the visual stimulus cueing the production of specific words). If two runs were performed, the first run was discarded and the second run was entered in the analysis. One participant required a second run of the category fluency task (see Supplemental Table 1).

### MRI Data Acquisition Parameters

Whole-brain BOLD imaging was conducted with an 8-channel head coil on a 3-Tesla GE Signa scanner located at the University of Rochester Medical Imaging Center. At the start of each participant’s scanning session, high-resolution structural T1 contrast images were acquired using a fast spoiled gradient echo pulse sequence (FSPGR BRAVO; TR = 108 ms, TE = 4ms, flip angle = 13°, FOV = 512mm, matrix = 512 × 512, 140 sagittal left-to-right slices, 1.3 × 0.48 × 0.48 mm per slice). An echo-planar imaging pulse sequence was used for T2* contrast (TR = 3000ms, TE = 35ms, flip angle = 90°, FOV = 256 × 256 mm, matrix = 64 × 64, 34 inferior-to-superior axial slices, voxel size = 3.75 × 3.75 × 4mm). Seventy volumes of BOLD data were collected (210 seconds per task).

Neurotypical participant MRI data were acquired using a 3-Tesla Siemens MAGNETOM Trio scanner with a 32-channel head coil at the University of Rochester. High-resolution structural T1 contrast images were acquired using a magnetization-prepared rapid gradient echo pulse sequence at the start of each participant’s scanning session (TR = 2530, TE = 3.44ms, flip angle = 7°, FOV = 256mm, matrix = 256 × 256, 192 1 x 1 x 1mm sagittal left-to-right slices). Diffusion MRI was acquired using a single-shot echo-planar sequence (60 diffusion directions with b = 1000 s/mm^2^, 10 images with b = 0 s/mm^2^, TR = 8900 msec, TE = 86 msec, FOV = 256 x 256mm, matrix = 128 x 128, voxel size = 2 x 2 x 2 mm, 70 inferior-to-superior axial slices; anterior-to-posterior phase encoding). A double-echo gradient echo field map sequence (echo time difference = 2.46 msec, EPI dwell time = 0.75 msec) was acquired with the same resolution as the DTI sequence to correct for distortion caused by B0 inhomogeneity.

### Lesion Identification

FSL’s ‘FLIRT’ function was used to set the voxels of each participant’s T1 anatomical image to 1 x 1 x 1 mm isotropic (140 sagittal left-to-right slices). Lesions were then identified by segmenting healthy tissue from damaged tissue visible on the native T1 anatomy. Lesioned voxels consisting of both grey and white matter were assigned a value of 1, and preserved voxels were assigned a value of 0 using ITK-SNAP software (Yushkevich et al., 2006). Lesion drawings were inspected by a neuro-radiologist naïve to the aims of the project (co-author S.P. Meyers). The neuro-radiologist provided study team members with corrective feedback when the drawings did not accurately capture the boundaries of the lesion. Lesion drawings in native T1 anatomical space were then entered as cost function masks (Brett et al., 2001) during anatomical and functional data pre-processing within fMRIPrep (Esteban et al., 2019). This feature ensures that the anatomical data could be normalized to Montreal Neurological Institute (MNI) space while minimizing error introduced by the lesion. The corrected lesion drawings were then normalized to MNI space using the transformation matrix that normalized the native T1 anatomy into MNI space using the ANTS toolbox (Avants et al., 2011).

### Anatomical Data Pre-processing

Each participant’s T1-weighted (T1w) images was corrected for intensity non-uniformity with ‘N4BiasFieldCorrection’ (ANTs), and was used as the T1w-reference throughout the workflow. The T1w-reference was then skull-stripped with the ‘antsBrainExtraction.sh’ workflow (from ANTs, within NiPype), using OASIS30ANTs as target template. Brain tissue segmentation of cerebrospinal fluid, white-matter, and gray-matter was performed on the brain-extracted T1w using ‘fast’ (FSL). Volume-based spatial normalization to standard space (MNI152nLin2009cAsym) was performed through nonlinear registration with ‘antsRegistration’, using brain extracted versions of both T1w reference and the T1w template. The ICBM 152 Nonlinear Asymmetrical template version 2009c was selected for spatial normalization.

### Functional Data Pre-processing

For each run of fMRI data, a reference volume and its skull-stripped version were generated using fMRIPrep. Head-motion parameters with respect to the BOLD reference (transformation matrices, and six corresponding rotation and translation parameters) were estimated before spatiotemporal filtering using ‘MCFLIRT’ from FSL. BOLD runs were slice-time corrected using ‘3dTshift’ from AFNI. The BOLD time-series were then resampled onto their original, native space by applying the transforms to correct for head-motion. The BOLD reference was then co-registered to the T1w reference using ‘mri_coreg’ (FreeSurfer) followed by ‘FLIRT’ (FSL) with the boundary-based registration cost-function. Co-registration was configured with twelve degrees of freedom to account for distortions remaining in the BOLD reference. The BOLD time-series were resampled into standard space, generating a preprocessed BOLD run in MNI152nLin2009cAsym space. All resamplings can be performed with a single interpolation step by composing all the pertinent transformations (i.e., head-motion transform matrices, susceptibility distortion correction when available, and co-registrations to anatomical and output spaces). Gridded (volumetric) resamplings were performed using ‘antsApplyTransforms’ (ANTs), configured with Lanczos interpolation to minimize the smoothing effects of other kernels.

Following pre-processing in fMRIPrep, fMRI and T1 anatomical data were analyzed with the BrainVoyager software package (Version 22.4) and in-house scripts drawing on the NeuroElf toolbox written in MATLAB. All functional data underwent temporal high-pass filtering (cutoff: 2 cycles per time course within each run), were smoothed at 6 mm FWHM, and were interpolated to 3 x 3 x 3 mm voxels. A participant-specific general linear model (GLM) was used to fit beta estimates to the experimental events of interest. Experimental events were convolved with a standard 2-gamma hemodynamic response function. The first derivatives of 3D motion correction from each run were added to all models as regressors of no interest to attract variance attributable to head movement (XYZ translation and rotation; 6 covariates). Finally, a group GLM was created to inspect the brain areas exhibiting differentially greater BOLD responses when engaging in the category and letter fluency tasks.

### Defining the Left Pars Triangularis and the Left Pars Opercularis

We used the left pars triangularis and the left pars opercularis ROIs from the Harvard-Oxford Cortical Probabilistic Atlas to extract contrast-weighted *t*-values using the contrast of ‘Fluency > Fixation’ separately in the category and letter fluency task. The voxelwise mean of all contrast-weighted *t*-values in each ROI served as the independent variable in subsequent voxel-based lesion activity mapping analyses.

### Voxel-based Lesion Activity Mapping (VLAM)

VLAM uses neural responses in an ROI to predict variance in voxelwise lesion incidence throughout the brain. VLAM was performed using the SVR-LSM toolbox (DeMarco & Turkeltaub, 2018). Only voxels lesioned in at least 10% of participants were included. We controlled for variability in lesion volume using ‘Direct Total Lesion Volume Control’, which divides lesioned voxels in each participant’s lesion map by the square root of the total lesion volume (mm^3^). This correction method places a greater emphasis on smaller lesions when controlling for each participant’s lesion volume. We investigated task-specific responses by entering contrast weighted *t*-values from the second fluency task as a covariate (e.g., to control for responses in the letter fluency task when evaluating the responses from the category task, and vice versa). Four VLAM analyses were conducted: two using neural responses in the left pars triangularis, and two using neural responses in the left pars opercularis.

For each analysis, voxelwise statistical significance was determined using a permutation analysis, whereby neural responses were randomly assigned to a lesion map, and the same procedure as described above was iterated 10,000 times. Voxelwise z-scores were computed for the true data in relation to the mean and standard deviation of voxelwise null distributions; the resulting z-score map was set to a threshold of z < -1.65 (*p* < .05, one-tailed) to determine chance-level likelihood of identifying a significant effect in each voxel. If the resulting clusters did not survive permutation analysis, we removed voxels that did not form a cluster of at least 500 contiguous voxels (for precedent, see (Burns et al., 2025; Garcea & Buxbaum, 2023; Garcea et al., 2020; Garcea et al., 2019; Garcea et al., in press; Lacey et al., 2017; Skipper-Kallal et al., 2017). The Harvard-Oxford Cortical Probabilistic Atlas was used to assess the location of significant voxels.

### Lesion Sites Common to Category and Letter Fluency

To determine the lesion sites common to both tasks, we re-ran each VLAM analysis without entering the neural responses from the second fluency task as a covariate. For each analysis, we identified the voxels that met the dual criteria of 1) Exhibiting an above-threshold z-score (z < -1.65), and 2) Forming a cluster of at least 500 contiguous voxels.

### Deterministic Tractography in Neurotypical Participants

FSL’s BET tool skull-stripped each participant’s diffusion and T1 images, as well as the fieldmap magnitude image. The B0 image was stripped from the diffusion weighted image, and the fieldmap was prepared using FSL’s fieldmap tool. Smoothing and regularization was performed using FSL’s fugue tool and a 3D Gaussian smoothing was applied (sigma = 4 mm). The magnitude image was warped based on this smoothing, with *y* as the warp direction. Eddy current correction was performed using FSL’s eddy_correct tool, which takes each volume of the diffusion-weighted image and registers it to the b0 image to correct for both eddy currents and motion. The deformed magnitude image was registered to the b0 image using FSL’s FLIRT tool. The resulting transformation matrix was then applied to the prepared fieldmap. Lastly, the diffusion-weighted image was undistorted using the registered fieldmap with FSL’s fugue tool. Intensity correction was also applied to this unwarping.

Diffusion data were analyzed with the DSI_Studio program. First, we created a database modeling fractional anisotropy (FA) along each voxel of the 54 neurotypical participants’ pre-processed diffusion data. Within DSI_Studio, the accuracy of b-table orientation was examined by comparing fiber orientations with those of a population-averaged template (Yeh et al., 2018). Tensor metrics were calculated using a b-value lower than 1750 s/mm². A deterministic fiber tracking algorithm (Yeh et al., 2013) was used with augmented tracking strategies (Yeh et al., 2021) to improve reproducibility. The anisotropy threshold was randomly selected between 0.5 and 0.7 otsu threshold. The angular threshold was randomly selected from 45 degrees to 90 degrees. The step size was set to voxel spacing. Tracks with length shorter than 30.0 or longer than 200.0 mm were discarded.

Next, we used the left pars triangularis as a seed ROI along with the voxels identified in each VLAM analysis. VLAM-identified voxels were treated as a second ROI to identify streamlines that pass through both the pars triangularis and the distal ROI. For each analysis, 1,000,000 seeds were placed to identify streamlines. Topology-informed pruning (Yeh et al., 2019) was applied to the tractography with 4 iteration(s) to remove false positive connections. Identical analyses using the left pars opercularis as the ROI and voxels identified in the VLAM analyses of the category and letter fluency tasks then followed, for a total of 4 analyses.

## Results

### Whole-brain analysis of category fluency and letter fluency

A whole-brain contrast map was generated for each fluency task using a group-level GLM. Engaging in the category and letter fluency tasks elicited robust BOLD responses in bilateral inferior, middle, and superior frontal gyri, bilateral insula, bilateral precentral gyri, bilateral intraparietal sulci bordering on the superior parietal lobule, and in bilateral posterior inferior and middle temporal gyri (see Supplemental Figure 1 for each group-level contrast map).

### Correlation of neural responses in the left pars triangularis and left pars opercularis

There were robust correlations between the neural responses in the category and letter fluency tasks in left pars triangularis (r(50) = 0.58, *p* <.001; see Figure 2A) and the left pars opercularis (r(50) = 0.52, *p* <.001; see Figure 2B and Supplemental Table 1). Thus, prior to conducting the category fluency VLAM analysis, neural responses from the letter fluency task were entered as a covariate to isolate the lesion sites uniquely associated with reduced neural responses in the category fluency task (and vice versa for the letter fluency VLAM analysis).

**Figure 2.**
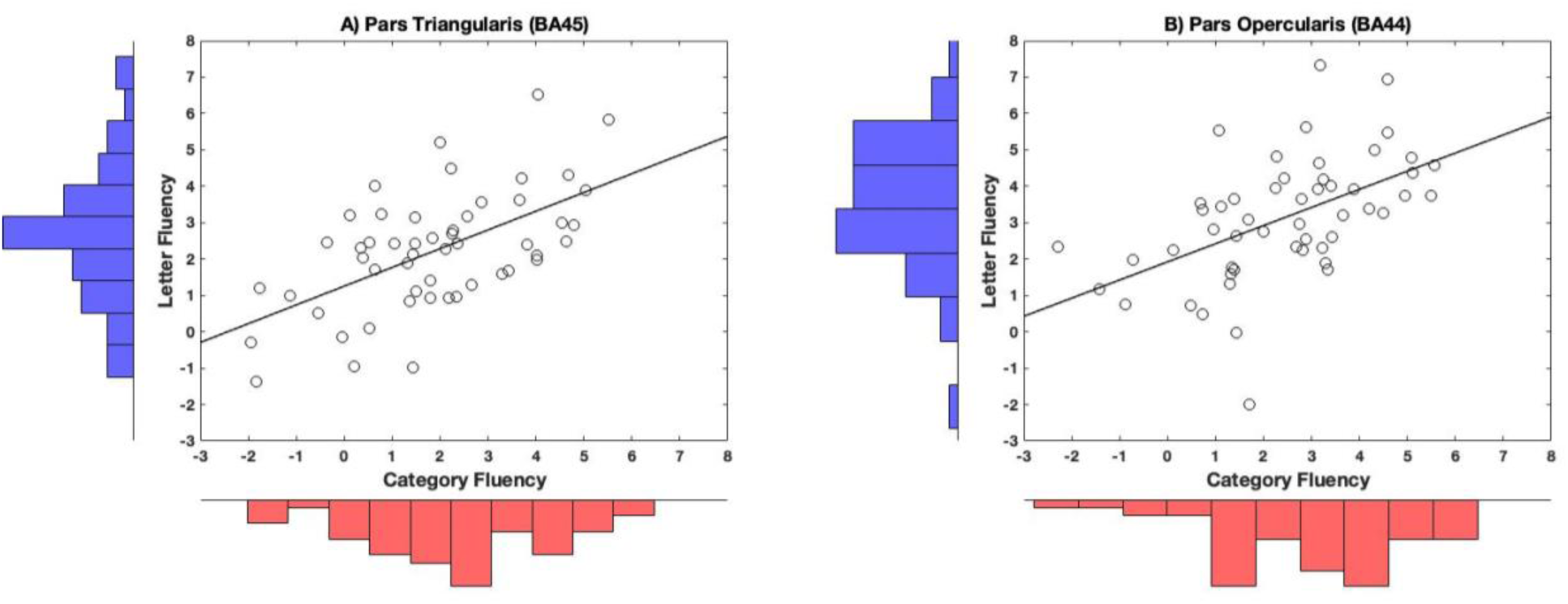
The relation between neural responses across the letter and category fluency tasks within each left inferior frontal gyrus ROI. There was a robust correlations between neural responses in category and letter fluency tasks in the left pars triangularis (r(50) = 0.58, *p* < .001; panel A) and the left pars opercularis (r(50) = 0.52, *p* < .001; panel B). A Shapiro-Wilk test confirmed that each distribution was normally distributed.

### Voxel-based Lesion-Activity Mapping Results

#### Category Fluency

The first hypothesis tested was that participants with lesions to anterior portions of the left middle and superior temporal gyri will show reduced neural responses in the left inferior frontal gyrus during category fluency (controlling for neural responses during letter fluency). When using the left pars triangularis as the reference ROI, we found that that lesion presence in the left temporal pole, anterior superior temporal gyrus, hippocampus, and amygdala was inversely related to the amplitude of neural responses in the left pars triangularis. This cluster included the left insula, the left hippocampus, the anterior division of the left parahippocampal gyrus, the left orbitofrontal cortex, the left planum polare, left amygdala, and the left putamen (see Figure 3A and Table 1A). When using the left pars opercularis as the reference ROI, we did not find evidence to support this hypothesis. We found that lesions to the right hippocampus and amygdala were inversely associated with neural responses in the left pars opercularis (Figure 4A and Table 2A). When removing the letter fluency covariate from the model, we found that lesions to the left temporal pole, anterior superior temporal gyrus, hippocampus, and amygdala were associated with reduced neural responses in pars opercularis (see Supplemental Figure 3). This finding replicates the VLAM result observed using the left pars triangularis ROI (Figure 2A).

**Figure 3.**
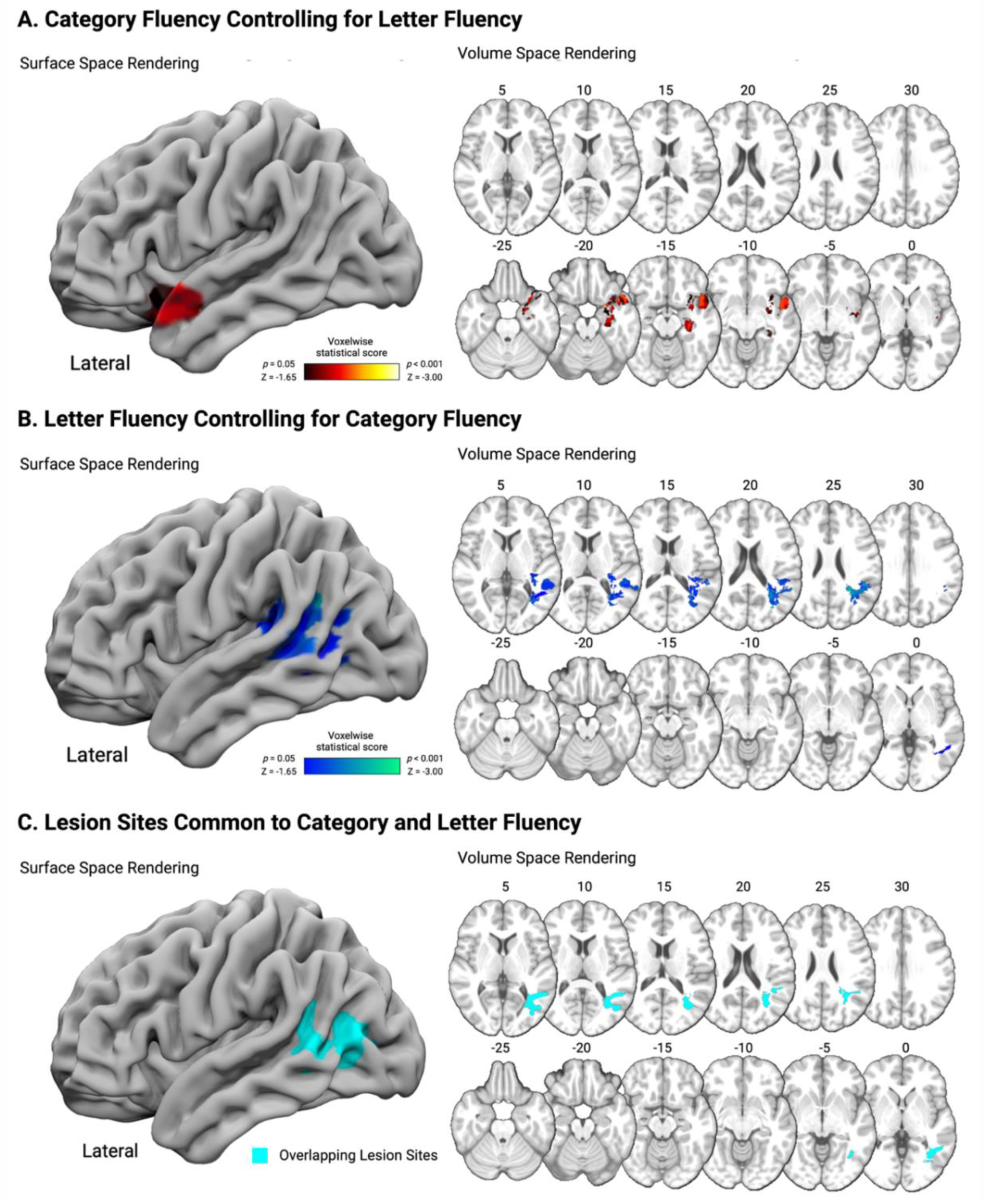
VLAM analysis of category and letter fluency using in the left pars triangularis ROI. Surface-based (left) and volumetric (right; MNI Z coordinate listed above axial slices, in radiological convention) renderings of lesion sites associated with decreased neural responses during category fluency controlling for letter fluency (A) and letter fluency controlling for category fluency (B). Lesion sites common to category fluency and letter fluency are rendered in cyan (C).

**Table 1.** Left hemisphere regions and MNI coordinates of voxels identified in the VLAM analysis of (A) category (B) letter fluency in the left pars triangularis. . The number of significant voxels in each identified region from the Harvard-Oxford cortical and subcortical parcellation atlases are provided, and the percentage of significant voxels in each atlas region is listed in parentheses (minimum threshold of 1%). The minimum threshold was relaxed to show the voxels common to both analyses in panel C.

| Region Name | Region Number | Peak MNI Coordinate (XYZ) |  |  | Statistical Value | Number of Significant Voxels |
| --- | --- | --- | --- | --- | --- | --- |
| A. Category Fluency Controlling for Letter Fluency |  |  |  |  |  |  |
| Hippocampus | 9 | -33 | -16 | -16 | -2.77 | 1378 (23.45%) |
| Temporal Pole | 15 | -24 | 7 | -26 | -2.58 | 2231 (10.95%) |
| Orbitofrontal cortex | 65 | -25 | 8 | -26 | -2.58 | 558 (3.92%) |
| Insular cortex | 3 | -33 | 6 | -16 | -2.29 | 646 (5.63%) |
| Planum Polare | 87 | -47 | 3 | -9 | -2.28 | 200 (6.57%) |
| Amygdala | 10 | -23 | 0 | -21 | -2.26 | 451 (17.76%) |
| Parahippocampal Gyrus, Anterior Division | 67 | -22 | 0 | -27 | -2.18 | 108 (2.22%) |
| Superior Temporal Gyrus, Anterior Division | 17 | -51 | 1 | -14 | -2.03 | 30 (1.28%) |
| Putamen | 6 | -28 | -1 | -8 | -1.99 | 90 (1.33%) |
| B. Letter Fluency Controlling for Category Fluency |  |  |  |  |  |  |
| Supramarginal Gyrus, Posterior Division | 39 | -50 | -49 | 30 | -2.66 | 497 (5.39%) |
| Parietal Operculum | 85 | -47 | -40 | 26 | -2.66 | 509 (11.12%) |
| Angular Gyrus | 41 | -43 | -54 | 27 | -2.57 | 288 (3.23%) |
| Planum Temporale | 91 | -53 | -36 | 13 | -2.47 | 443 (9.64%) |
| Middle Temporal Gyrus, Temporooccipital Division | 25 | -53 | -49 | 8 | -2.39 | 542 (7.56%) |
| Superior Temporal Gyrus, Posterior Division | 19 | -54 | -36 | 5 | -2.02 | 42 (1.23%) |
| C. Overlapping Lesion Sites |  |  |  |  |  |  |
| Middle Temporal Gyrus, Temporooccipital Division | 25 | -52 | -51 | 7 | - | 625 (8.72%) |
| Lateral Occipital Cortex, Inferior Division | 45 | -43 | -69 | 8 | - | 1007 (6.06%) |
| Lateral Occipital Cortex, Superior Division | 43 | -40 | -69 | 16 | - | 156 (0.37%) |
| Inferior Temporal Gyrus, Temporooccipital Division | 31 | -42 | -57 | -4 | - | 8 (0.14%) |
| Supramarginal Gyrus, Posterior Division | 39 | -51 | -48 | 22 | - | 6 (0.07%) |
| Parietal Operculum | 85 | -39 | -40 | 19 | - | 1 (0.02%) |
| Planum Temporale | 91 | -53 | -43 | 20 | - | 1 (0.02%) |

**Figure 4.**
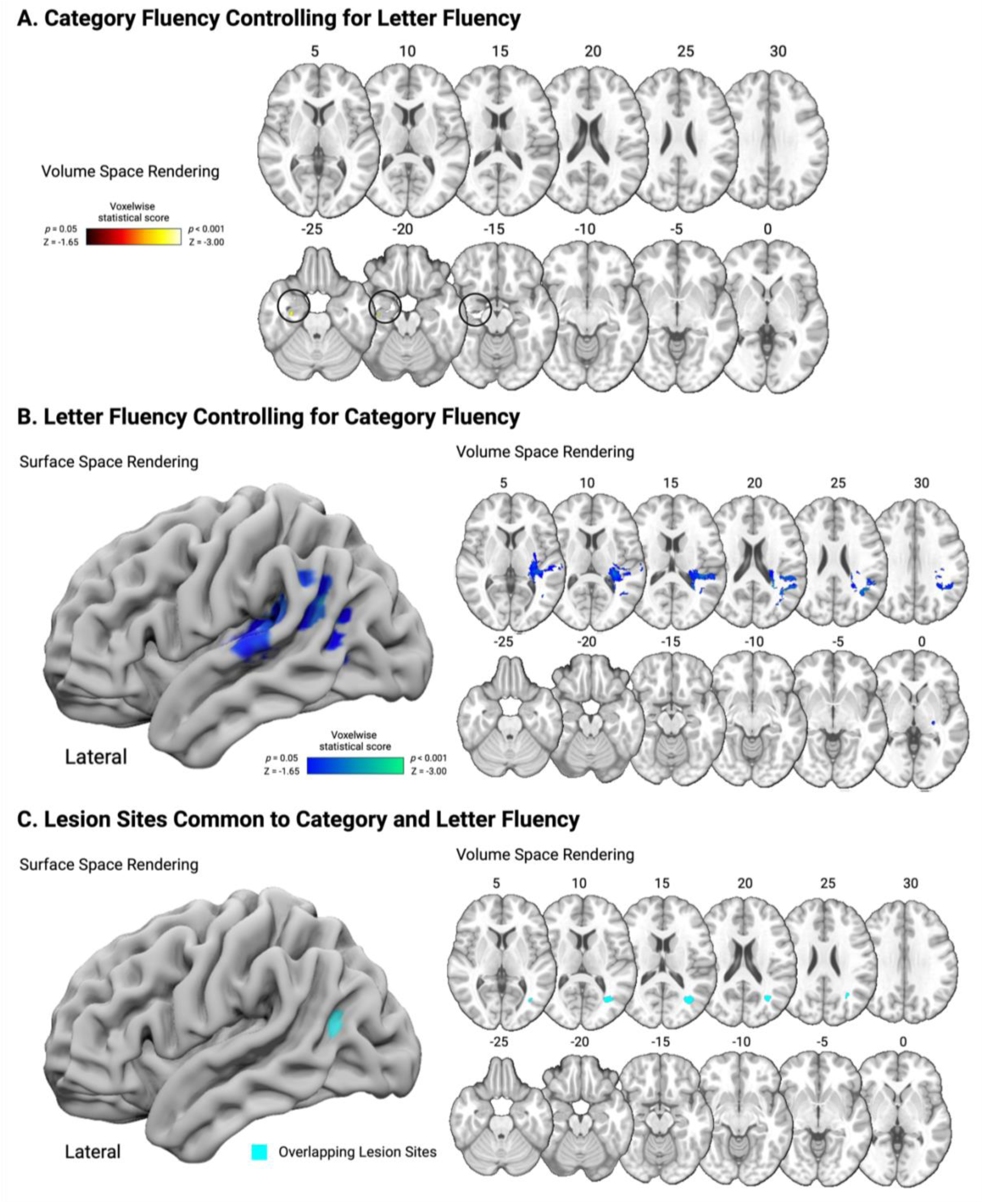
VLAM analysis of category and letter fluency using the left pars opercularis ROI. Surface-based (left) and volumetric (right; MNI Z coordinate listed above axial slices, in radiological convention) renderings of lesion sites associated with decreased neural responses during category fluency controlling for letter fluency (A; volume space only; see circled voxels) and letter fluency controlling for category fluency (B). Lesion sites common to category fluency and letter fluency are rendered in cyan (C).

**Table 2.**
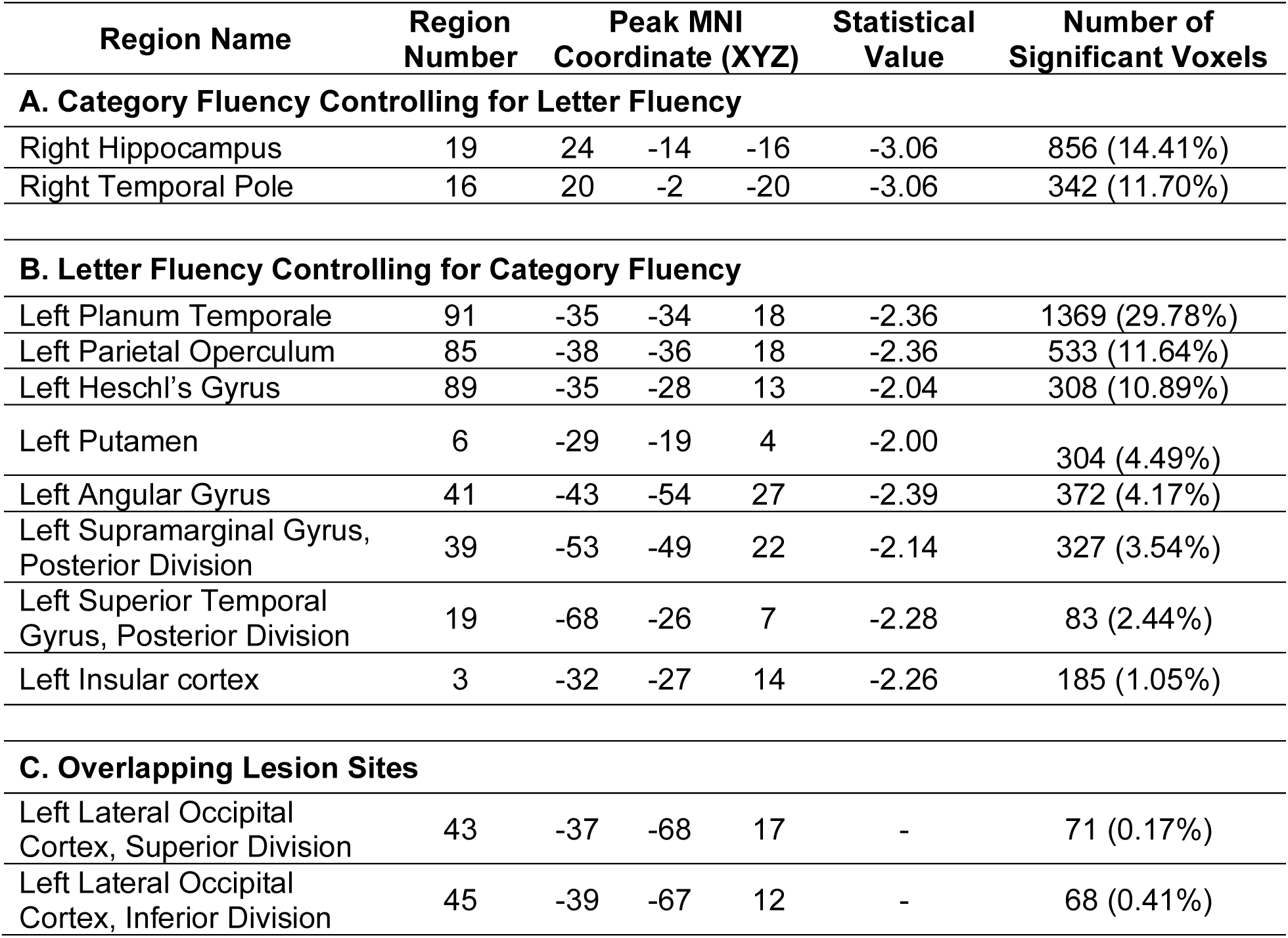
Left and right hemisphere regions and MNI coordinates of voxels identified in the VLAM analysis of (A) category (B) letter fluency in the left pars opercularis. The number of significant voxels in each identified region from the Harvard-Oxford cortical and subcortical parcellation atlases are provided, and the percentage of significant voxels in each atlas region is listed in parentheses (minimum threshold of 1%). The minimum threshold was relaxed to show the voxels common to both analyses in panel C.

| Region Name | Region Number | Peak MNI Coordinate (XYZ) |  |  | Statistical Value | Number of Significant Voxels |
| --- | --- | --- | --- | --- | --- | --- |
| A. Category Fluency Controlling for Letter Fluency |  |  |  |  |  |  |
| Right Hippocampus | 19 | 24 | -14 | -16 | -3.06 | 856 (14.41%) |
| Right Temporal Pole | 16 | 20 | -2 | -20 | -3.06 | 342 (11.70%) |
| B. Letter Fluency Controlling for Category Fluency |  |  |  |  |  |  |
| Left Planum Temporale | 91 | -35 | -34 | 18 | -2.36 | 1369 (29.78%) |
| Left Parietal Operculum | 85 | -38 | -36 | 18 | -2.36 | 533 (11.64%) |
| Left Heschl's Gyrus | 89 | -35 | -28 | 13 | -2.04 | 308 (10.89%) |
| Left Putamen | 6 | -29 | -19 | 4 | -2.00 | 304 (4.49%) |
| Left Angular Gyrus | 41 | -43 | -54 | 27 | -2.39 | 372 (4.17%) |
| Left Supramarginal Gyrus, Posterior Division | 39 | -53 | -49 | 22 | -2.14 | 327 (3.54%) |
| Left Superior Temporal Gyrus, Posterior Division | 19 | -68 | -26 | 7 | -2.28 | 83 (2.44%) |
| Left Insular cortex | 3 | -32 | -27 | 14 | -2.26 | 185 (1.05%) |
| C. Overlapping Lesion Sites |  |  |  |  |  |  |
| Left Lateral Occipital Cortex, Superior Division | 43 | -37 | -68 | 17 | - | 71 (0.17%) |
| Left Lateral Occipital Cortex, Inferior Division | 45 | -39 | -67 | 12 | - | 68 (0.41%) |

#### Letter Fluency

The second hypothesis tested was that participants with lesions to the left posterior superior temporal gyrus and supramarginal gyrus will exhibit reduced neural responses in the left inferior frontal gyrus during letter fluency. We found evidence in support of this hypothesis using both inferior frontal gyrus ROIs. Lesion presence in the left planum temporale, the left parietal operculum, posterior superior temporal gyrus, posterior portion of the left supramarginal gyrus, and the left angular gyrus was inversely related to the strength of neural responses in the left pars triangularis (see Figure 3B and Table 1B). This lesion site included the temporooccipital division of the left middle temporal gyrus, and the superior and inferior divisions of the left lateral occipital cortex. A similar pattern was identified using the left pars opercularis as the ROI. Specifically, lesions to the left planum temporale, posterior superior temporal gyrus, parietal operculum, posterior supramarginal gyrus, and angular gyrus were inversely related to the strength of neural responses during letter fluency but not category fluency (see Figure 4B and Table 2B). This lesion site included the left putamen and left insular cortex.

#### Overlapping Lesion Sites

Each VLAM analysis was re-run without entering neural responses from the other fluency task as a covariate (see Supplemental Figures 2 and 3). In the analysis of the left pars triangularis, overlapping voxels were identified in the white matter undercutting the left posterior supramarginal gyrus, parietal operculum, and planum temporale, and in the left posterior middle and inferior temporal gyri and the left posterior lateral occipital cortex (see Figure 3C and Table 1C). In the analysis of the left pars opercularis, a small portion of the white matter undercutting the left posterior lateral occipital cortex was identified (see Figure 4C and Table 2C).

### Deterministic tractography in neurotypical participants

#### Category Fluency

The voxels in Figures 3A and 3B were entered as seed ROIs to identify the white matter tracts connecting those VLAM-identified lesion sites with the left pars triangularis in neurotypical participants. The same analysis was carried out using the voxels in Figure 4A and 4B and the left pars opercularis. We hypothesized that the anterior temporal lesion site identified in the VLAM analysis of category fluency would be structurally connected to the left inferior frontal gyrus via the left IFOF and/or the left uncinate fasciculus. We find evidence supporting this hypothesis: 84.82% of the fibers identified were in the left IFOF (blue), while a smaller portion of fibers identified included the left extreme capsule (9.28%; yellow), the left arcuate fasciculus (3.58%; red), and the left uncinate fasciculus (2.32%; green; see Figure 5A and Table 5A). When using the left pars opercularis and the right hippocampal lesion site identified in the VLAM analysis of category fluency (Figure 4A), no white matter fibers were identified.

**Figure 5.**
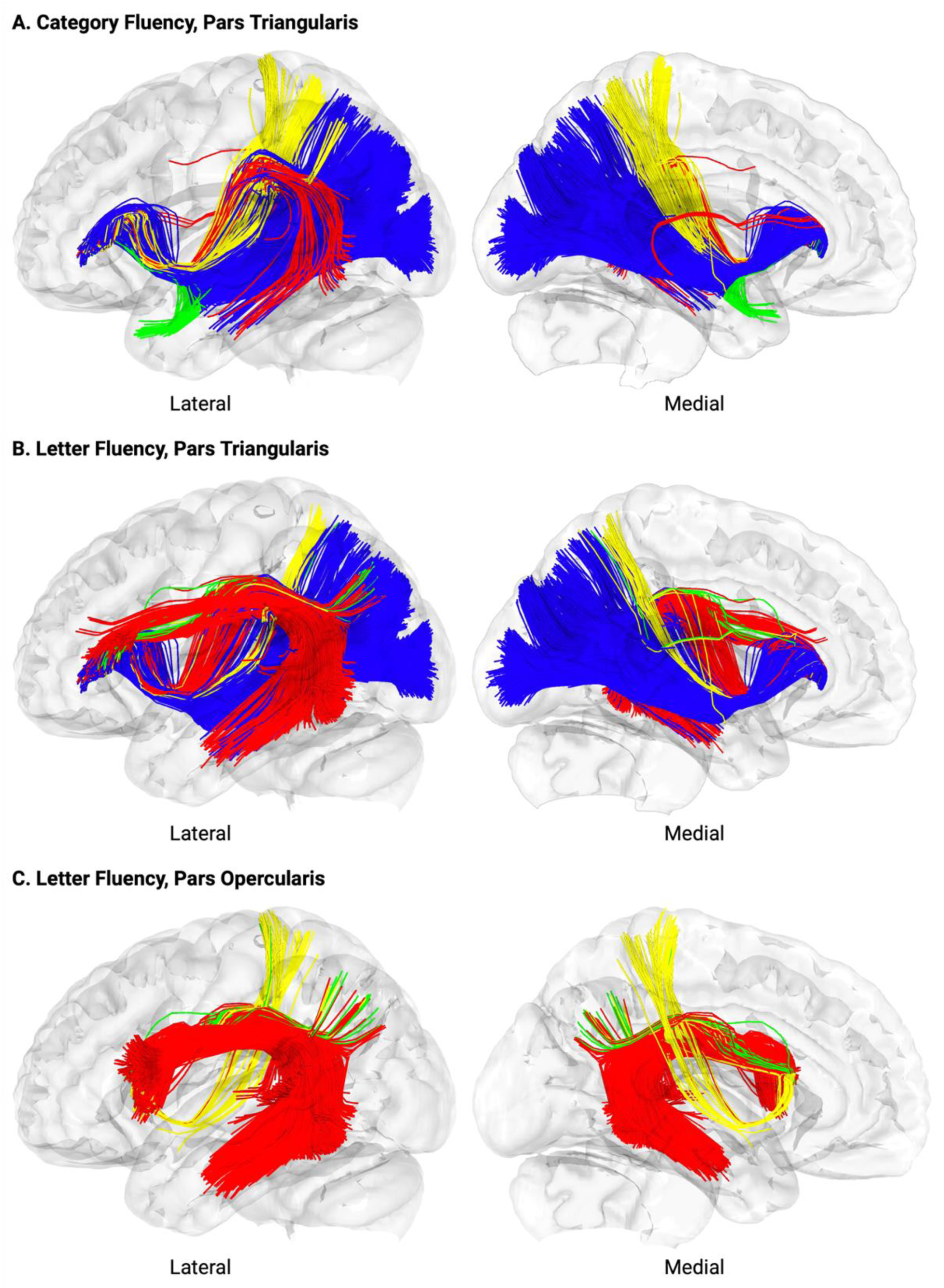
White matter fibers connecting the VLAM-identified lesion sites to the left pars triangularis. (A) The left IFOF (blue) is identified as the principal white matter pathway connecting the left pars triangularis to the left anterior temporal lobe lesion sites from the category fluency VLAM analysis. (B) Though the left IFOF (blue) was identified again as the principal white matter pathway, the left arcuate fasciculus (red) was also identified as a major white matter pathway connectivity the left pars triangularis to the left temporo-parietal lesion sites found in the VLAM analysis of letter fluency. (C) When inspecting the white matter connecting the left pars opercularis and the temporo-parietal lesions sites identified in the letter fluency VLAM analysis, the left arcuate fasciculus (red) was identified as the principal white matter pathway.

**Table 3.** Left hemisphere fiber tracts identified in the deterministic tractography analysis.

| Fiber tract name | Number of fibers identified and the proportion of fibers within each tract |
| --- | --- |
| **A. White matter fibers connecting the left pars triangularis to lesion sites identified in the VLAM analysis of category fluency.** |  |
| Inferior Fronto-Occipital Fasciculus | 1827 (84.82%) |
| Extreme Capsule | 200 (9.28%) |
| Arcuate Fasciculus | 77 (3.58%) |
| Uncinate Fasciculus | 50 (2.32%) |
| **B. White matter fibers connecting the left pars triangularis to lesion sites identified in the VLAM analysis of letter fluency.** |  |
| Inferior Fronto-Occipital Fasciculus | 1611 (70.94%) |
| Arcuate Fasciculus | 578 (25.45%) |
| Extreme Capsule | 63 (2.77%) |
| Second branch of the Superior Longitudinal Fasciculus | 19 (0.84%) |
| **C. White matter fibers connecting the left pars opercularis to lesion sites identified in the VLAM analysis of letter fluency.** |  |
| Arcuate Fasciculus | 1915 (95%) |
| Extreme Capsule | 78 (3.87%) |
| Second branch of the Superior Longitudinal Fasciculus | 22 (1.13%) |

#### Letter Fluency

We hypothesized that the lesion site identified in the VLAM analysis of letter fluency would be structurally connected to the left inferior frontal gyrus via the left inferior longitudinal fasciculus and/or the left arcuate fasciculus. Across both analyses we find support for this hypothesis, with graded differences as a function of the inferior frontal ROI. When using the left pars triangularis, 70.94% of the fibers were in the left IFOF (blue), 25.45% of fibers were in the left arcuate fasciculus (red), 2.77% included the extreme capsule (yellow), and less than 1% of the fibers were from second branch of the left superior longitudinal fasciculus (see Figure 5B and Table 5B). When using the left pars opercularis, 95% of the fibers were in the left arcuate fasciculus (red); the remaining 5% included left extreme capsule (yellow; 3.87%) and the second branch of the left superior longitudinal fasciculus (green; 1.13%; see Figure 5C and Table 5C).

### General Discussion

Activation in the left inferior frontal gyrus represents a summation of endogenous control processes that guide the search of task-relevant representations. In the context of the verbal fluency task, that search occurs over semantic or phonologically related concepts. Thus, the goal of this investigation was to combine fMRI with Voxel-based Lesion-Activity Mapping (VLAM) to test how lesions to anterior temporal and temporo-parietal areas that represent semantic or phonological information would dampen neural responses in the left inferior frontal gyrus in a task-specific manner. We first showed that lesion damage to the left anterior temporal lobe and hippocampus were inversely related to the strength of neural responses in the left pars triangularis specifically during the category fluency task (Figure 3A). This effect was not observed in the left pars opercularis; instead, we found that lesions to the right hippocampus were associated with reduced neural responses during category fluency (Figure 4A). By contrast, left temporo-parietal lesions were inversely related to the strength of neural responses in the left pars triangularis (Figure 3B) and the left pars opercularis (Figure 4B) specifically during the letter fluency task.

Although common lesion sites and white matter fibers were found across the tasks, we identified partially distinct white matter tracts connecting the VLAM-derived lesion sites to the left inferior frontal gyrus. Across deterministic tractography analyses, the left IFOF emerged as the principal structural pathway connecting the left pars triangularis to the lesion sites from the category and letter fluency VLAM analyses (Figure 5A and 5B). The left arcuate fasciculus was identified as a second white matter tract connecting the left pars triangularis to the lesions identified in the letter fluency VLAM analysis (Figure 5B). By contrast, the left arcuate fasciculus was the principal structural pathway connecting the left pars opercularis to the lesion sites from the letter fluency VLAM analysis (Figure 5C). These results suggest that anatomically dissociable brain networks interact with the left inferior frontal gyrus when different search strategies constrain the retrieval of conceptual representations for word production.

### VLAM Results

Our VLAM analysis of category fluency identified left anterior temporal lobe regions previously found in behavioral VLSM investigations (e.g., see Baldo et al., 2006; Biesbroek et al., 2016; Zigiotto et al., 2022). These findings suggest the left pars triangularis forms a network together with the anterior temporal lobe and hippocampus, among other regions, to shape the retrieval of semantically related concepts. In the same participants, we show that lesions to these anterior temporal sites do not lead to attenuated neural responses in the left pars triangularis during the letter fluency task. VLSM studies of aphasic patients have shown that lesions to the anterior temporal lobe can lead to semantic errors in picture naming (Baldo et al., 2013; Cloutman et al., 2009; Mirman, Zhang, et al., 2015; Walker et al., 2011). Our findings, coupled with these prior observations, provide converging evidence that one key function of the left anterior temporal lobe is to support differentiation among semantically related concepts for further lexical-semantic processing (e.g., see Schwartz et al., 2009).

When entering the letter fluency task as a covariate, lesions to left anterior temporal regions were not associated with reduced neural responses in the left pars opercularis during category fluency (Figure 4A). However, when removing the covariate from the model, we did find that left anterior superior temporal gyrus lesions were associated with reduced neural responses in the left pars opercularis (see Supplemental Figure 3). One possibility is that neural responses in the left pars opercularis reflect domain-general retrieval, articulatory, or cognitive control processes shared across both verbal fluency tasks. Under this interpretation, entering letter fluency as a covariate may remove variance associated with these shared processes. With respect to the left pars triangularis, the persistence of the VLAM category fluency effect in the anterior temporal lobe after accounting for letter fluency is consistent with prior proposals that anterior portions of the left inferior frontal gyrus contribute more selectively to retrieval of semantically related concepts (e.g., see Katzev et al., 2013; Poldrack et al., 1999).

By contrast, across both pars triangularis and pars opercularis, reduced neural responses during letter fluency (but not category fluency) were associated with lesions to left supramarginal gyrus, parietal operculum, and superior temporal gyrus. These findings demonstrate that a distinct network supports the retrieval of phonologically related concepts for word production. Interestingly, there is overlap between the left temporo-parietal lesion sites we identified and prior investigations of the lesion sites associated with phonological errors in picture naming (Mirman, Chen, et al., 2015; Schwartz et al., 2012), and reduced word production during letter fluency (e.g., see Blecher et al., 2019; Chouiter et al., 2016), with a common lesion locus in the left supramarginal gyrus. These observations are consistent with dual-stream neurocognitive models which propose that the dorsal stream is crucial for phonological sequencing, articulatory planning, and sensorimotor integration during word production (for discussion, see Dell et al., 2013; Fridriksson et al., 2018; Hickok & Poeppel, 2004; Schwartz, 2014).

A key open question is whether functional interactions among frontal, temporal, and inferior parietal regions are bidirectional. We have shown that damage to the search of semantically or phonologically related concepts in anterior temporal and posterior temporo-parietal cortex, respectively, can shape endogenous control processes in the inferior frontal cortex. Because we did not have adequate lesion coverage in the frontal lobe, we are unable to test if lesions to the left pars triangularis or opercularis disrupt neural processing in distal temporo-parietal regions in a task-specific manner. One study by Price and colleagues (2001) showed that lesions to the left inferior frontal gyrus led to reduced fMRI responses in the left inferior temporal cortex when reading written words. Interestingly, those lesions did not disrupt neural responses for written word reading in the left middle temporal gyrus. These results demonstrate that task-specific disruptions in neural processing from frontal lesions can occur, a phenomenon Price and colleagues refer to as dynamic diaschisis (for discussion, see Carrera & Tononi, 2014). Future studies are needed to investigate if lesions to distinct inferior frontal gyrus regions disrupt neural responses in the anterior temporal lobe specifically in the category fluency task but not in the letter fluency task, and vice versa for neural responses in temporo-parietal cortex in the letter fluency task. This work will be crucial to advance understanding of the directionality of network interactions underlying the endogenous selection of conceptual representations for word production.

### Deterministic Tractography Results

The left IFOF was identified as the principal tract connecting each VLAM-derived lesion site with the left pars triangularis. The IFOF, together with the inferior longitudinal fasciculus, are two white matter tracts that constitute the ventral language pathway (Dick et al., 2014). Prior studies in awake neurosurgery patients have shown that semantic paraphasias in picture naming can be elicited from direct electrical stimulation to the left IFOF (Duffau et al., 2005; Duffau et al., 2008). A small portion of the extreme capsule fiber pathway was also identified in both deterministic fiber tracking analyses. The extreme capsule connects middle portions of the middle and superior temporal gyri with anterior portions of the inferior frontal gyrus (e.g., see Dick et al., 2014; Makris & Pandya, 2009) and is implicated in semantic analysis for language comprehension (Saur et al., 2008) and word production in the category fluency task (Barbeau et al., 2024). Our results add to this prior literature by showing that the left IFOF and left extreme capsule are two ventral-going tracts that integrate semantic representations in the temporal lobe with endogenous control processes in the frontal lobe in support of word production in the verbal fluency task.

In contrast to those ventral stream tracts, the left arcuate fasciculus was identified when using the VLAM-derived lesion sites from the letter fluency task. This was found when using both left pars triangularis, and, to a greater extent, when using the left pars opercularis (see Figure 5C). The left arcuate fasciculus is the principal white matter tract that makes up the dorsal stream of language (Hickok & Poeppel, 2004; Ivanova et al., 2021; Saur et al., 2008). Prior studies have shown that reduced structural integrity of the left arcuate fasciculus was associated with fewer words produced in the letter fluency task but not the category fluency task (e.g., see Blecher et al., 2019). This finding does not imply that the dorsal stream supports word production during letter fluency but not category fluency. Our proposal is that word production in the letter fluency task emphasizes a phonologically-constrained search strategy, such as syllabification of the initial letter, which drives neural processing in the left posterior superior temporal gyrus and left supramarginal gyrus, two regions that are connected to the left pars opercularis (and to a lesser extent the left pars triangularis) via the left arcuate fasciculus.

### Future Directions and Conclusions

Word production in category and letter fluency may rely on a common search strategy. For example, participants may search for phonologically similar words from a common semantic category (e.g., foods that start with the letter S; Schwartz et al., 2003). Though a cognitive neuroscientist reviewed the task with every participant before the scan to ensure task compliance, audio recordings were not acquired during the scan. Thus, we are unable to validate whether participants were using such a strategy. Nevertheless, common lesion sites were identified, including the white matter undercutting the left posterior middle temporal gyrus and the inferior division of the left lateral occipital cortex (see Figure 3C and Figure 4C). This lesion site converges with what has been referred to as a white matter “bottleneck” in the posterior temporal lobe (Griffis et al., 2017; see also Mirman, Chen, et al., 2015 for discussion of white matter bottlenecks)—a small area with numerous crossing white matter fibers, including the left arcuate fasciculus and left IFOF (Fernandez-Miranda et al., 2015). One possibility is that a small amount of damage to these crossing fibers may structurally disconnect the left inferior frontal gyrus from the left posterior temporal lobe, leading to disruptions in neural processing as has been observed here.

In summary, we use neural responses in the left inferior frontal gyrus to test whether distinct temporal lobe areas constrain the retrieval of semantically or phonologically related words. Using VLAM and deterministic fiber tracking, we show that functional interactions between the left pars triangularis and left anterior temporal lobe, mediated via the IFOF and extreme capsule, underlie the retrieval of words in the category fluency task. The retrieval of words in the letter fluency task drives interactions in a distinct network, including the left pars triangularis and opercularis, and the left temporo-parietal cortex, connected via the IFOF, extreme capsule, and arcuate fasciculus. These data support a model whereby functional interactions among structurally connected networks facilitate the retrieval of concepts for lexical selection and subsequent word production.

## Data availability

The whole-brain maps of contrast-weighted t-values, lesions in MNI space, and analysis code will be made available via a publicly accessible repository upon publication of the manuscript. Participants did not consent to public archival of the MRI data. Researchers interested in accessing the raw MRI data should contact the corresponding author to establish a data use agreement.

## Acknowledgements

We thank Alena Stasenko and Mary Abbe Roe for their assistance de-identifying the MRI DICOM files.

## SUPPLEMENTAL ONLINE MATERIALS

**Supplemental Table 1.**
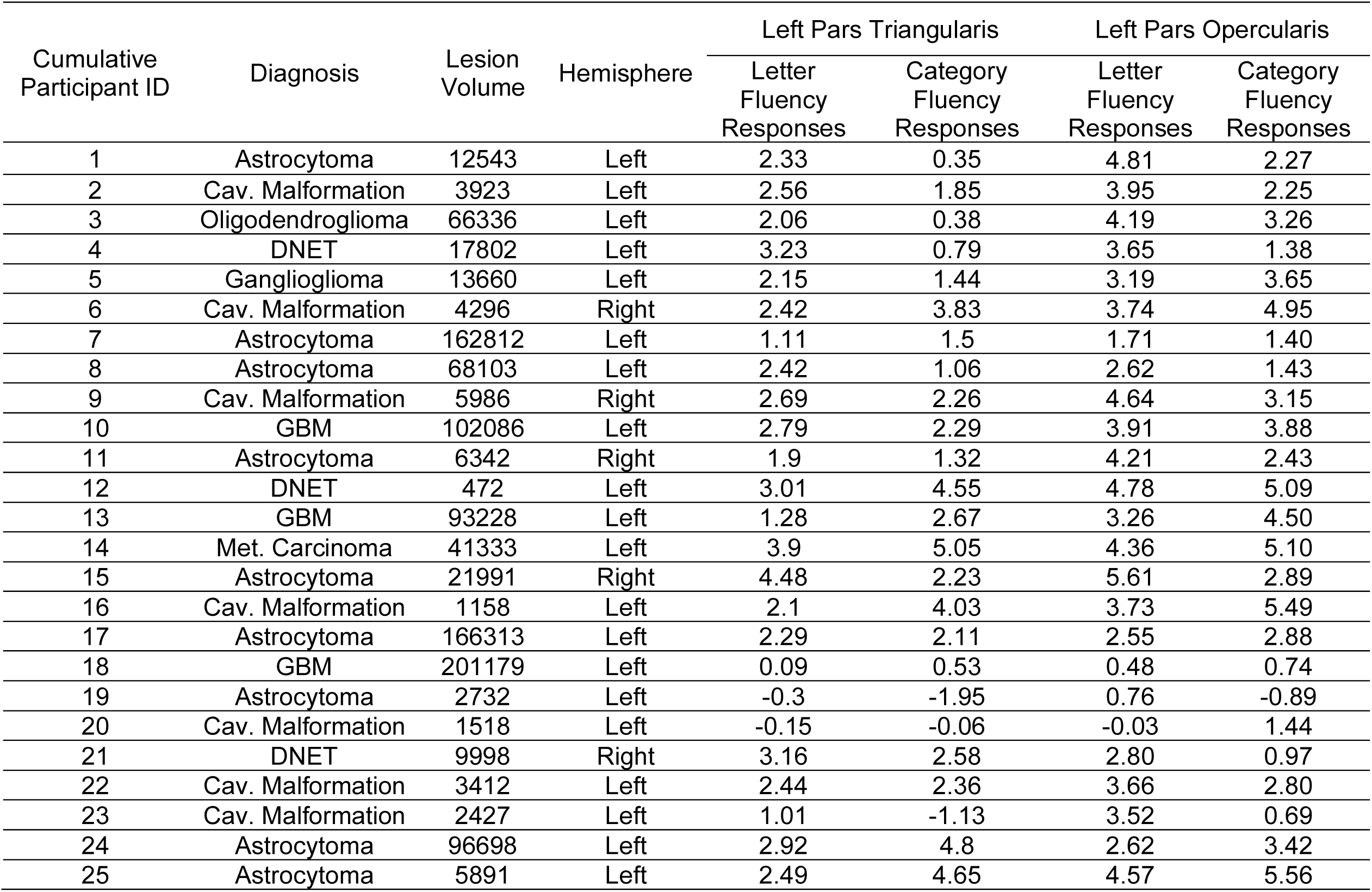
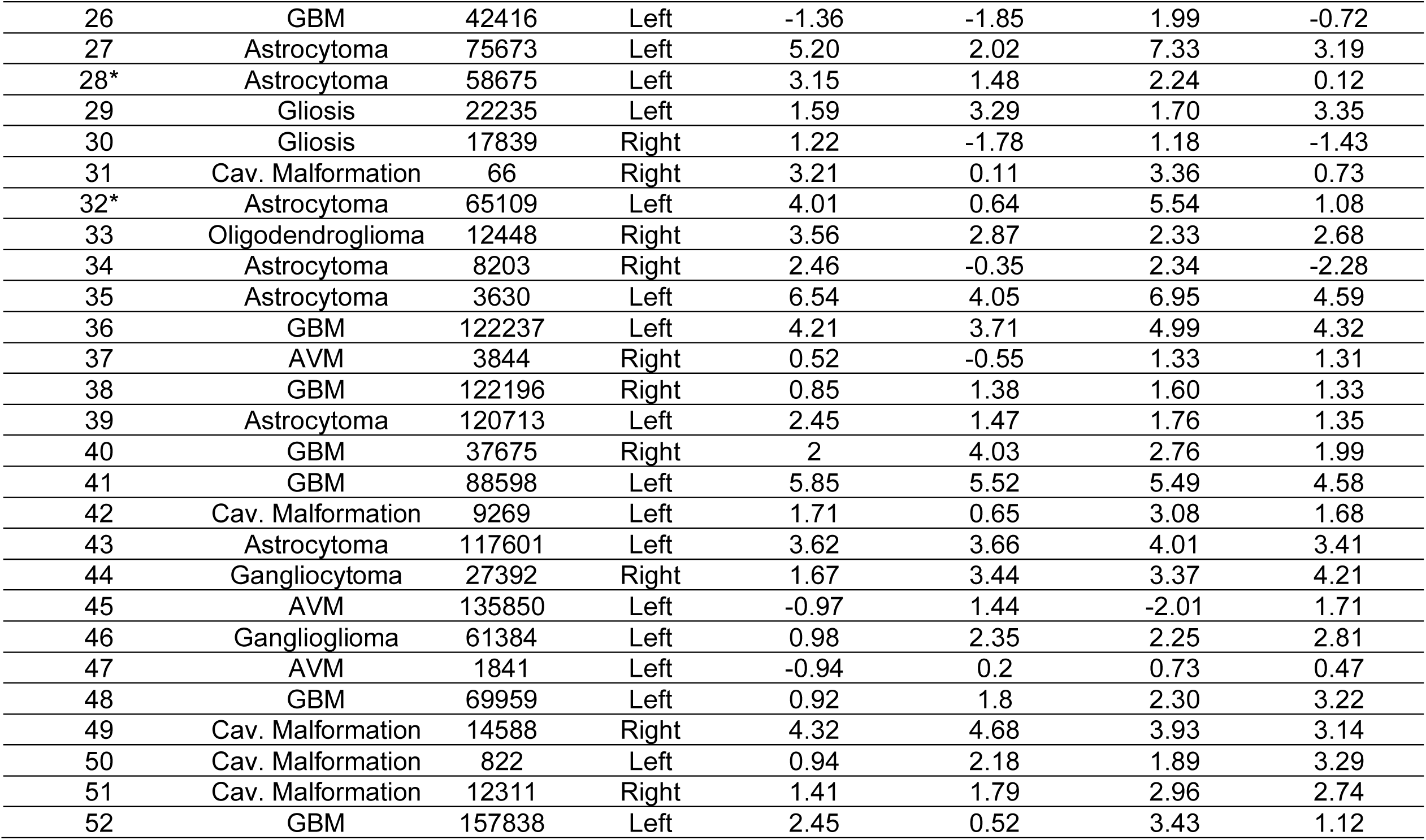
Demographic information for each participant, along with contrast-weighted *t*-values for each task and ROI. *Abbreviations.* AVM, Arteriovenous Malformation; Cav. Malformation, Cavernous Malformation; DNET, Dysembryoplastic Neuroepithelial Tumor; GBM, Glioblastoma Multiforme; Met. Carcinoma, Metastatic Carcinoma. * Participant completed a second run of the category fluency task.

**Supplemental Figure 1.**
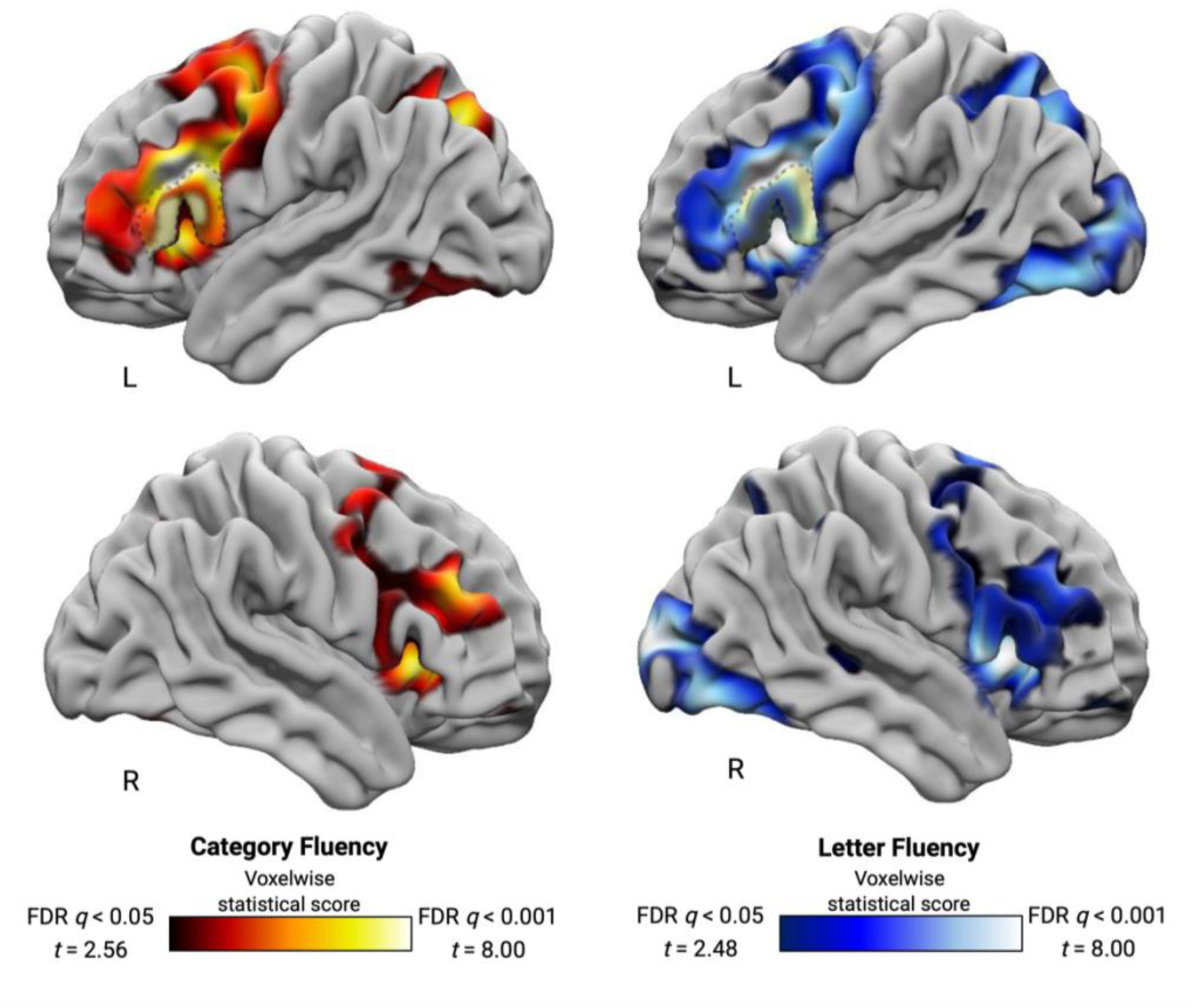
Whole-brain contrast of [‘Fluency > Fixation’] for the Category Fluency Task (left) and the Letter Fluency Task (right). Each map was FDR corrected (*q* < 0.05). Outlined on the lateral surface of the left hemisphere is a representation of the left pars triangularis and left pars opercularis. These ROIs were used to extract participant-specific contrast-weighted t-values for subsequent VLAM analyses.

**Supplemental Figure 2.**
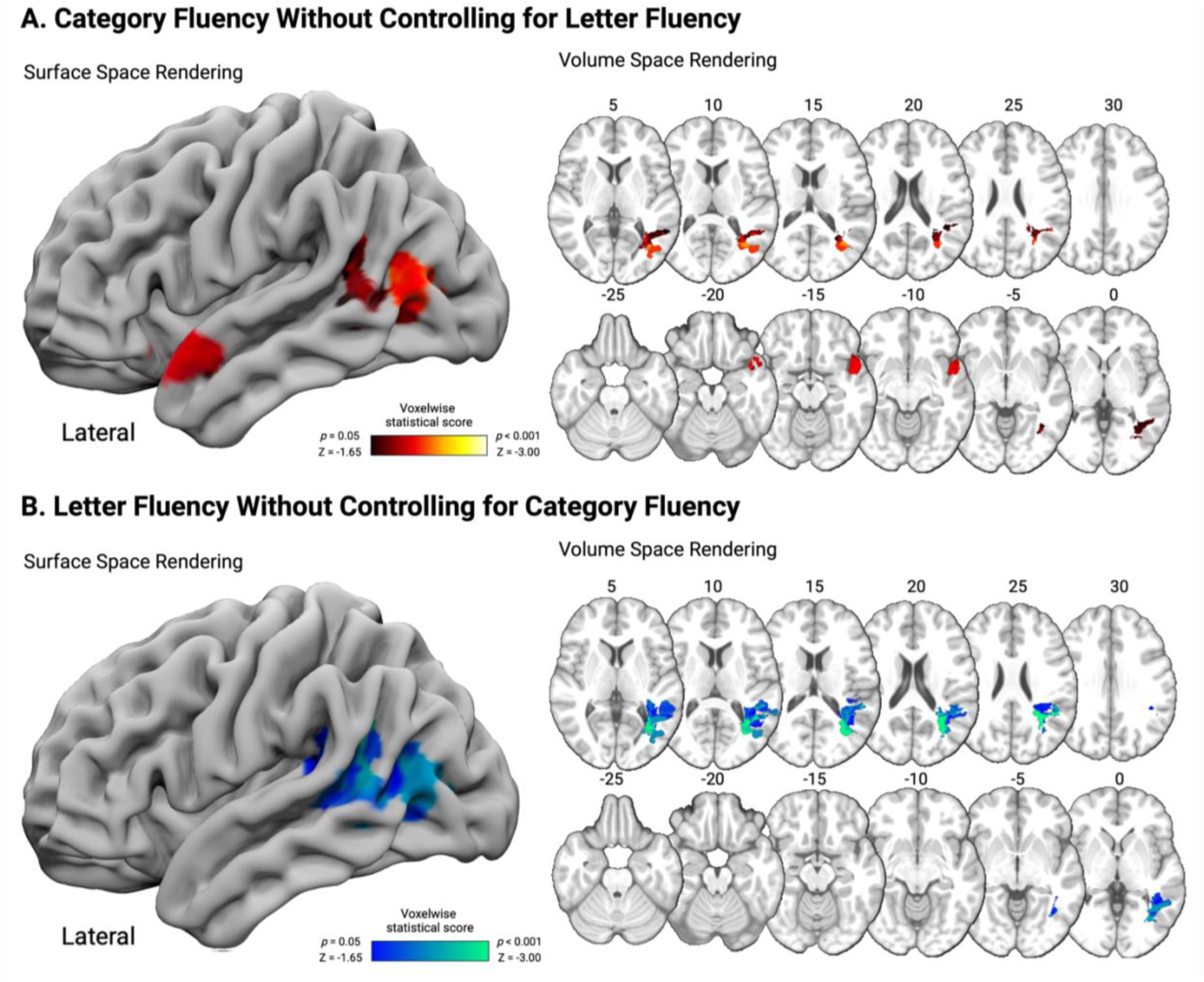
VLAM analysis when using the left pars triangularis as the reference ROI. The lesion sites of the category fluency VLAM analysis without controlling for neural responses in the letter fluency task are in panel A, while the lesion sites of the letter fluency VLAM analysis without controlling for neural responses in the category fluency task are in panel B.

**Supplemental Figure 3.**
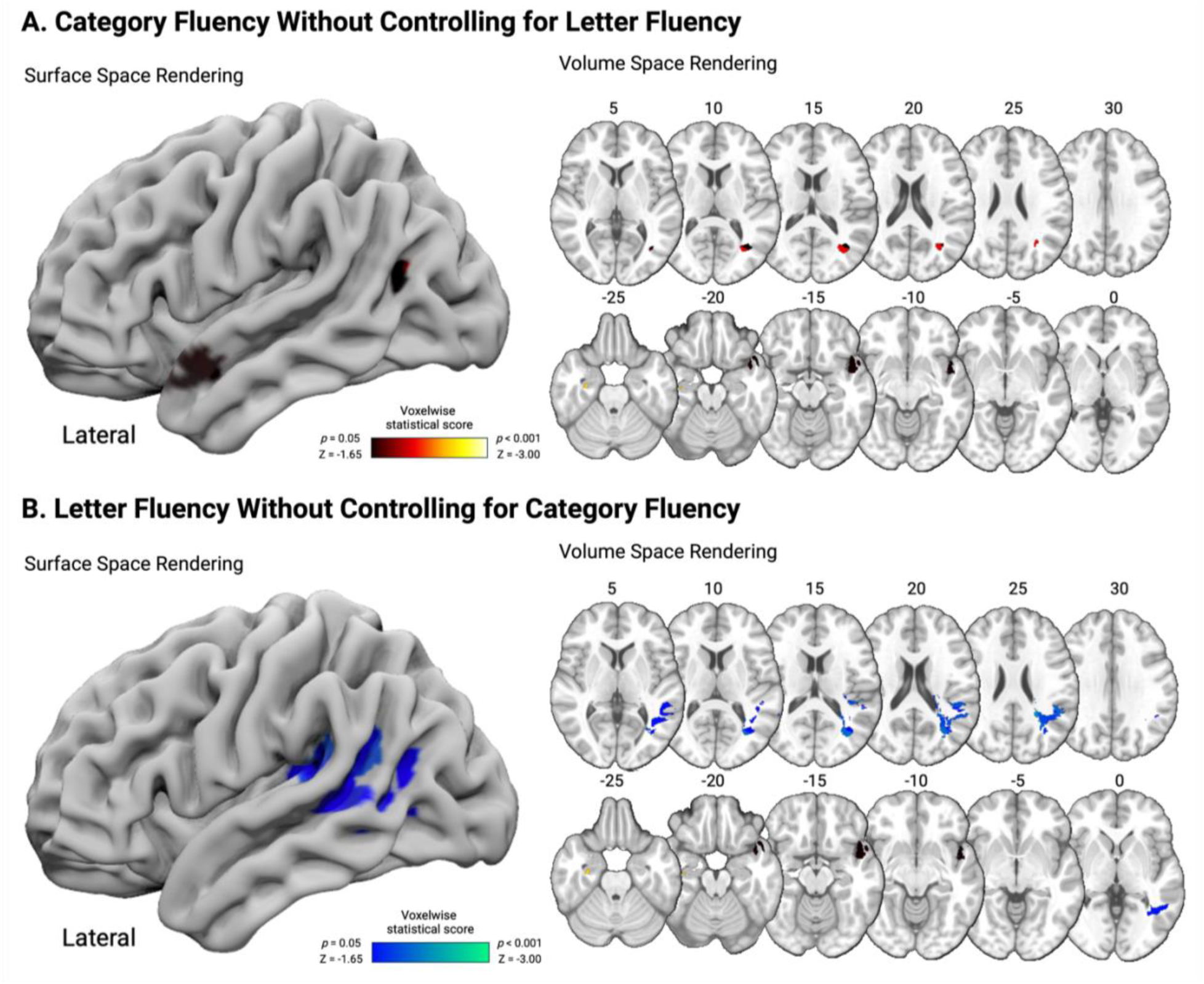
VLAM analysis when using the left pars opercularis as the reference ROI. The lesion sites of the category fluency VLAM analysis without controlling for neural responses in the letter fluency task are in panel A, while the lesion sites of the letter fluency VLAM analysis without controlling for neural responses in the category fluency task are in panel B.

